# Effect of Heterogeneity in Recombination Rate on Variation in Realised Relationship

**DOI:** 10.1101/341776

**Authors:** Ian M.S. White, William G. Hill

## Abstract

Individuals of specified pedigree relationship vary in the proportion of the genome they share identical by descent, i.e. in their realised or actual relationship. Basing predictions of the variance in realised relationship solely on the proportion of the map length shared implicitly assumes that both recombination rate and genetic information are uniformly distributed along the genome, ignoring the possible existence of recombination hotspots, and failing to distinguish between coding and non-coding sequences. In this paper we quantify the effects of heterogeneity in recombination rate at broad and fine scale levels on the variation in realised relationship. A chromosome with variable recombination rate usually shows more variance in realised relationship than does one having the same map length with constant recombination rate, especially if recombination rates are higher towards chromosome ends. Reductions in variance can also be found, and the overall pattern of change is quite complex. In general, local (fine-scale) variation in recombination rate, e.g. hotspots, has a small influence on the variance in realised relationship. Differences in rates across longer regions and between chromosome ends can increase or decrease the variance in realised relationship, depending on the genomic architecture.

## INTRODUCTION

Pedigree relationship defines the probability of identity by descent of paternally and maternally derived sites at any point in the autosomal genome. Because of linkage, sites on the same chromosome do not segregate independently, causing pairs of individuals of specified pedigree relationship to vary in the proportion of the genome they share identical by descent (ibd), i.e. in their *actual* or *realised* relationship. For a parent and offspring, or for monozygotic twins, there is complete sharing and no variance, but for other relationships the variance is non-zero and depends on the number of chromosomes, on their map lengths, and on both the type and degree of the pedigree relationship (*R*), differing for example between that for half sibs and that for grandchild and grandparent, although *R* = 0.25 for both. With increasingly distant relatives, the variance of relationship falls but the CV increases. The analysis has been developed over a number of years (e.g. Franklin, 1977; Stam and Zeven, 1981; Hill, 1993; Guo, 1995), and a fairly comprehensive analysis for a broad range of relationships was given by Hill and Weir (2011). Visscher (2009) provides some approximate summary statistics for the human genome and suggests how the results can be utilised to estimate genetic variation in quantitative traits in non-pedigreed populations. Variation in realised inbreeding of offspring of related parents (Hill and Weir, 2011) and variation in relationship among partially inbred individuals (Hill and Weir, 2012) can be computed similarly.

Formulae and examples to date for the variance in realised relationship have been based almost exclusively on the proportion of the map length shared, thereby making two implicit but perhaps dubious biological assumptions. The first is that the recombination rate is uniform along the genome, although average recombination rates (expressed as centimorgans per megabase, cM/Mb, for example) typically differ among chromosomes. For example, they range from 0.96 to 2.11 in humans (Kong et al., 2004) and differ much more widely in birds, which typically have many small chromosomes with high and diverse recombination rates (Groenen et al., 2009). There are both broad scale and small scale differences within chromosomes, notably at recombination hotspots (McVean et al., 2004; Stapley et al., 2017). The second assumption is that the genetic information content is uniformly distributed, despite the presence of local differences, including noncoding regions such as introns.

Given the crucial role played by linkage disequilibrium in genome-wide association studies (GWAS), and in the genomic prediction of genotypic value of livestock and plants and of disease risk in humans, a better understanding of the effect of diversity in recombination rate is needed.

The magnitude of the variation in recombination rate is well recognised. Stapley et al. (2017) reviewed contributing factors, and Ritz et al. (2017) discussed the extent to which variation in recombination rate is itself adaptive. Since the PRDM9 gene was identified, analysis and discussion have been published on its evolution, its role in determining sequence-wide hot spots, and (possibly) speciation events (Schwartz et al., 2014). Factors that determine observed chromosome numbers, average recombination rates and distribution over chromosomes have been discussed in the evolutionary literature. It is likely there are many factors involved, including, for example, epistasis and reproductive isolation. Charlesworth and Charlesworth (2010, pp. 546–561) provide an extensive discussion.

There remain many questions as to why magnitudes and patterns of recombination rate take the values they do. In this paper we do not attempt to address these philosophical questions. Instead, as a basis for understanding, we quantify the effects of heterogeneity in recombination rate at broad and fine scale levels on the variation in realised relationship for a wide range of pedigrees. We develop methods to do so, finding a broken stick model particularly useful.

Observations of genomic identity between chromosomes at the molecular level are initially likely to be in terms of the physical length, measured in megabases (Mb), rather than in map lengths, measured in morgans (M). Most or all published calculations on prediction of the magnitude of genome sharing are at the level of map distance, however. The conversion from one to the other then depends on the correspondence of the physical and linkage maps. This varies among chromosomes and species around the typical mammalian figure of 1 cM/Mb, depending *inter alia* on positions of centromeres and repetitive regions. For example, the chicken has a very high cM/Mb ratio compared to mammals and indeed compared to the zebra finch (Backström et al., 2010), but for both species of birds the recombination rates on the microchromosomes are relatively high (Stapley et al., 2017). For human chromosomes, although the linkage map is not far from linear with respect to the physical map for the longer metacentric chromosomes, it is somewhat sigmoidal; whereas for the shortest acrocentric chromosome over 25% of the centromeric end shows no recombination (Matise et al., 2007).

Provided a chromosome has constant recombination rate, physical and map distance are equivalent, but with heterogeneity in the recombination rate, this is no longer the case. As results of Hill and Weir (2011) and others are presented in terms of map distances, there is potentially a need to convert physical lengths to map lengths before calculating variance of ibd sharing. We investigate whether their formulae are sufficiently robust to make such conversion unnecessary. Specifically, we compute the variation in realised relationship as a joint function of the physical and linkage map and then consider how much this influences variance in realised relationship. As the inbreeding of offspring is proportional to the relationship of their parents, the analysis also applies directly to variation in realised inbreeding.

## ANALYSIS

First we re-express some methods and results of Hill and Weir (2011) in a simpler form as a basis for further analysis. Descriptions and abbreviations for the various pedigree relationships mentioned are given in Table 1.

**Table 1:**
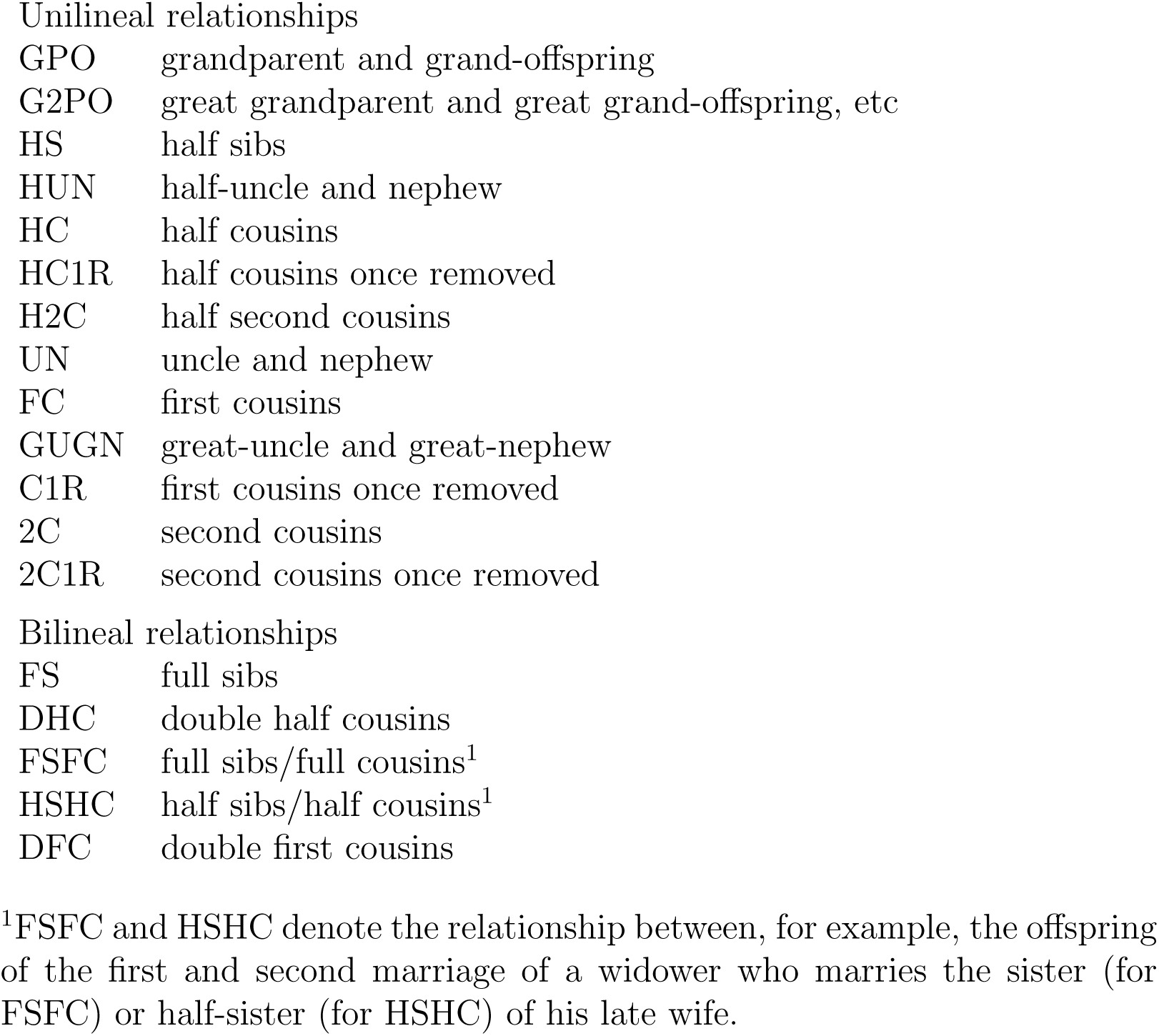
Relationships mentioned in tables, figures, and text

|  |  |
| --- | --- |
| GPO | grandparent and grand-offspring |
| G2PO | great grandparent and great grand-offspring, etc |
| HS | half sibs |
| HUN | half-uncle and nephew |
| HC | half cousins |
| HC1R | half cousins once removed |
| H2C | half second cousins |
| UN | uncle and nephew |
| FC | first cousins |
| GUGN | great-uncle and great-nephew |
| C1R | first cousins once removed |
| 2C | second cousins |
| 2C1R | second cousins once removed |
| Bilineal relationships |  |
| FS | full sibs |
| DHC | double half cousins |
| FSFC | full sibs/full cousins<sup>1</sup> |
| HS HC | half sibs/half cousins<sup>1</sup> |
| DFC | double first cousins |

<sup>1</sup>FSFC and HSHC denote the relationship between, for example, the offspring of the first and second marriage of a widower who marries the sister (for FSFC) or half-sister (for HSHC) of his late wife.

### Simplification and generalisation of basic formulae

We assume the genome to consist of a number of independently segregating chromosomes, and we focus on just one of these. Consider a pair of individuals, P and Q, who have a unilineal relationship, i.e. they are related on only one side of their pedigree. (See Table 2 for examples.) For such a relationship, it is possible to identify one chromosome in each individual through which the genetic relationship is expressed. Let *k* be the probability of ibd sharing at a single site on these two chromosomes. This is the probability that P and Q share exactly one pair of alleles ibd at the locus, denoted *k*_1_ by Hill and Weir (2011). Wright’s (1922) relationship coefficient is *R* = *k*/2, and *k*/4 is the coancestry coefficient for P and Q.

**Table 2:** Coefficients of the variance of identity required for eqn (1), for a range of relationships. The scale factor is *k*^2^ = (1/total)^2^.

| | $a_0$ | $a_1$ | $a_2$ | $a_3$ | $a_4$ | $a_5$ | $a_6$ | $a_7$ | $a_8$ | total |
| --- | --- | --- | --- | --- | --- | --- | --- | --- | --- | --- |
| Unilineal relationships |  |  |  |  |  |  |  |  |  |  |
| GPO | 1 | 1 | . | . | . | . | . | . | . | 2 |
| HS | 1 | . | 1 | . | . | . | . | . | . | 2 |
| UN | 1 | . | $\frac{1}{2}$ | $\frac{1}{2}$ | . | . | . | . | . | 2 |
| G2PO | 1 | 2 | 1 | . | . | . | . | . | . | 4 |
| HUN | 1 | 1 | 1 | 1 | . | . | . | . | . | 4 |
| GUGN | 1 | 1 | $\frac{1}{2}$ | 1 | $\frac{1}{2}$ | . | . | . | . | 4 |
| FC | 1 | . | $1\frac{1}{2}$ | 1 | $\frac{1}{2}$ | . | . | . | . | 4 |
| G3PO | 1 | 3 | 3 | 1 | . | . | . | . | . | 8 |
| HC | 1 | 2 | 2 | 2 | 1 | . | . | . | . | 8 |
| C1R | 1 | 1 | $1\frac{1}{2}$ | $2\frac{1}{2}$ | $1\frac{1}{2}$ | $\frac{1}{2}$ | . | . | . | 8 |
| G4PO | 1 | 4 | 6 | 4 | 1 | . | . | . | . | 16 |
| HC1R | 1 | 3 | 4 | 4 | 3 | 1 | . | . | . | 16 |
| 2C | 1 | 2 | $2\frac{1}{2}$ | 4 | 4 | 2 | $\frac{1}{2}$ | . | . | 16 |
| G5PO | 1 | 5 | 10 | 10 | 5 | 1 | . | . | . | 32 |
| 2C1R | 1 | 3 | $4\frac{1}{2}$ | $6\frac{1}{2}$ | 8 | 6 | $2\frac{1}{2}$ | $\frac{1}{2}$ | . | 32 |
| Bilineal relationships: double sharing |  |  |  |  |  |  |  |  |  |  |
| FS | 1 | . | 2 | . | 1 | . | . | . | . | 4 |
| FSFC | 1 | 0 | $2\frac{1}{2}$ | 1 | 2 | 1 | $\frac{1}{2}$ | 0 | 0 | 8 |
| HSHC | 1 | 2 | 3 | 4 | 3 | 2 | 1 | 0 | 0 | 16 |
| DFC | 1 | . | 3 | 2 | $3\frac{1}{4}$ | 3 | $2\frac{1}{2}$ | 1 | $\frac{1}{4}$ | 16 |
| DHC | 1 | 4 | 8 | 12 | 14 | 12 | 8 | 4 | 1 | 64 |

The amount of ibd sharing between P and Q is random because it is affected by recombination, and the variance of ibd sharing is related to the covariances of ibd status for all pairs of loci. For a range of different relationships, Hill and Weir (2011) calculate the probability of ibd sharing at two linked sites as a polynomial in 1 − *c,* where *c* is the recombination fraction. Here we follow the same approach, but express the probability as a polynomial in 1 − 2*c*, which leads to equivalent but simpler coefficients. Following Hill and Weir (2011), we assume that Haldane’s (1919) mapping function applies, so that 1 − 2*c* = exp(−2*d*), where *d* ≥ 0 is the map distance between sites. The probability of ibd sharing at these loci is then
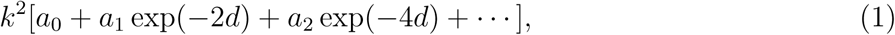

where *a*_0_, *a*_1_, etc are the polynomial coefficients, all of which are positive, or zero. Values of the coefficients for a range of relationships are given in Table 2. In the limit as *d* → ∞, when the two sites segregate independently, *a*_0_ = 1. Likewise, when *d* = 0 (the two sites segregate as one) the coefficients sum to 1*/k* = 2*^n^*. Dropping the first term gives the covariance of identity for two sites *d* morgans apart as

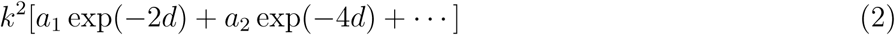

Expression (2) shows how, for a given relationship, covariance is attenuated by recombination as the map distance between sites increases.

The ibd process and an interpretation based on Markov chain theory is explained briefly in Appendix 1.

### Variation as a function of physical length

In order to compute the variance in relationship as a function of physical length, a natural (but not only possible) method is to weight each base in the DNA of the region equally. Hence, let *X* and *Y* denote physical positions on the chromosome, measured as proportions of total physical length, and let *ψ*(*X*) and *ψ*(*Y*) denote the corresponding map positions, expressed as proportions of total map length. Following Chakravarti (1991), we refer to *ψ* as the ‘Marey’ map, relating map length to physical length. (However, our axes are transposed compared to Chakravarti’s.) The derivative of *ψ* is the local recombination rate, i.e. the rate at which the proportion of map length increases with increasing proportion of physical length. Scaling by the ratio of map length of chromosome in cM to physical length of chromosome in Mb produces the conventional units cM/Mb. For a unilineal relationship, the variance of ibd for a chromosome of map length *ℓ*, *v*(*ℓ*), is obtained by averaging each term of expression (2) over all pairs of loci. Specifically, we replace *d* by *ℓ*(*ψ*(*X*) − *ψ*(*Y*)) and integrate exp[−2*sℓ*(*ψ*(*X*) − *ψ*(*Y*))] over the triangular region 0 *< Y* < *X* < 1 to obtain the average
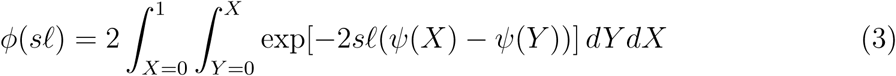

We assume that the number of genomic sites is large enough that the sum of terms can be approximated by (3). Then, from (2),
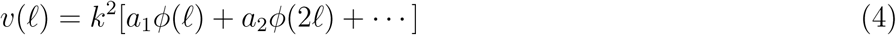

For the grandparent – grandoffspring relationship (GPO), *v*(*ℓ*) = *k*^2^*ϕ*(*ℓ*), and eqn (4) extends the result to other relationships, taking coefficients *a*_1_, *a*_2_, etc. from Table 2. For half-sibs (HS), for example, *v*(*ℓ*) = *k*^2^*ϕ*(2*ℓ*), so that the variance of sharing is the same for a HS relationship with map length *i* and a GPO relationship with map length 2*ℓ*.

For the special case of constant recombination rate (indicated by a zero suffix),
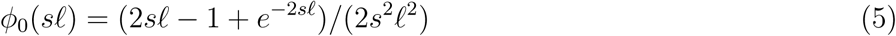

(Hill and Weir, 2011). The corresponding variance of relationship is

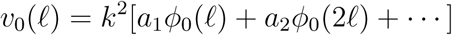

and the ratio *v*(*ℓ*)/*v*_0_(*ℓ*) is a measure of the effect of non-linearity in the Marey map which we refer to subsequently as the ‘discrepancy’ ratio, with values far from unity in either direction indicating a strong effect.

For bilineal relationships (e.g. full-sibs, double first cousins), where ibd sharing is possible through both paternal and maternal lineages, the overall proportion of ibd sharing is the average of two values, one from each lineage. Because the lineages segregate independently, the covariance of identity is the sum of two terms like (2), one for each lineage. With bilineal relationships, variation in ‘double’ sharing (simultaneous sharing in both lineages) can be calculated in a similar way to variance in single sharing, based on products of terms like (1) for each lineage. In the simplest cases, e.g. FS, DFC, when the chromosomal relationship is the same in both lineages, the bilineal variance is simply twice that for the corresponding unilineal relationship.

### Broken-stick models

A broken-stick model represents the relationship between physical and map distance on the chromosome as a continuous, piece-wise linear function. The recombination rate is assumed to be constant within defined chromosomal segments, but differs among segments. The variation in physical distance shared can then be calculated by summation, avoiding numerical integration.

Consider a single chromosome divided into segments, within each of which the recombination rate is constant. Denote the map length of the ith segment by *ℓ_i_*, and its physical length, expressed as a proportion of the total physical length of the chromosome, by *p_i_*, with 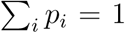. As an example, consider the case of grandparent and grand-offspring, for which the only non-zero coefficient in (4) is *a*_1_ = 1 (Table 2).

Let *S_i_* be the proportion of physical (and map) length shared in the ith segment. The calculation of var(*S_i_*) is the same as that for a whole chromosome with constant recombination rate: from (5),
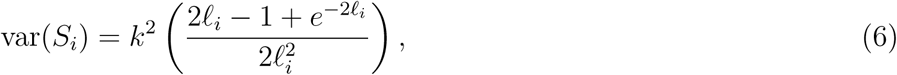

and the covariance of sharing between two segments is
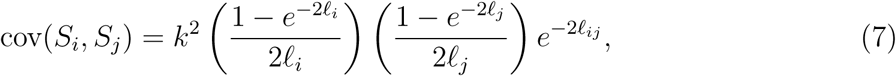

where *ℓ_ij_* is the total map length of segments lying between the ith and jth segments, with *ℓ_ij_* = 0 when they are adjacent.

For the whole chromosome, the proportion of physical length shared is 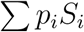, with variance
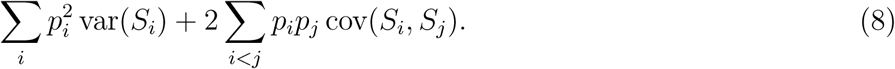

For other relationships in Table 2 the variance is calculated from (8), with eqn (6) modified to

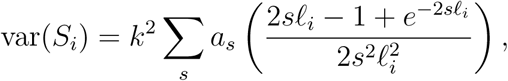

and with a similar modification to eqn (7).

### Data availability

The genomic data (Groenen et al., 2009, Supp. Table S1) analyzed below is available at https://genome.cshlp.org/content/19/3/510/suppl/DC1.

## RESULTS

### An illustrative example

Three examples of Marey maps are shown in Fig. 1. In one (afferent, ‘bringing inwards’), recombination is concentrated at the centre of the chromosome, and in the other (efferent, ‘conveying outwards’), there are high recombination rates near the chromosome ends. Half-way between these two extremes, the Marey map is linear, corresponding to a constant recombination rate.

**Figure 1:**
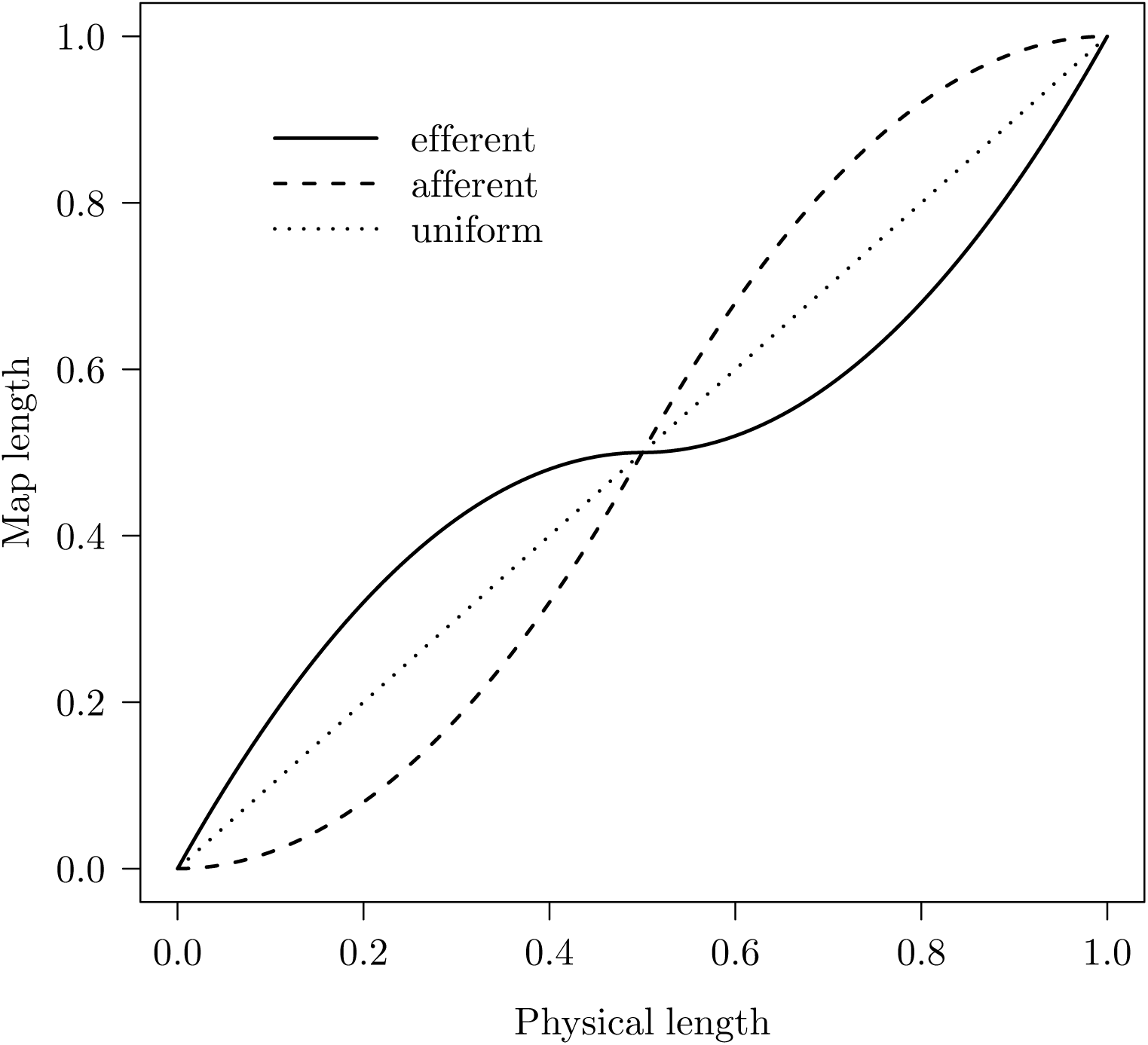
Three examples of Marey maps. Recombination is distributed away from the chromosome centre for the ‘efferent’ map, and concentrated at the centre for the ‘afferent’ map. Intermediate cases (not shown) produce weaker effects.

Consider the chromosome as divided into two segments of equal physical length, with *S*_1_ and *S*_2_ the proportions shared in each segment, so the proportion of the whole chromosome shared is (*S*_1_ + *S*_2_)/2, with variance *V*(1 + *ρ*)/2, where *V* = var(*S*_1_) = var(*S*_2_), and *ρ* is the correlation between *S*_1_ and *S*_2_. As the Marey map is changed from ‘efferent’ (recombination at ends) to ‘afferent’ (recombination at centre) *V* changes very little, but *ρ* steadily diminishes (Table 3). In this instance, the effect of a change in the Marey map (from efferent to afferent) is almost entirely the broad-scale effect on *ρ.* The fine-scale effect on *V* (the average pairwise covariance within segments) is negligible.

**Table 3:**
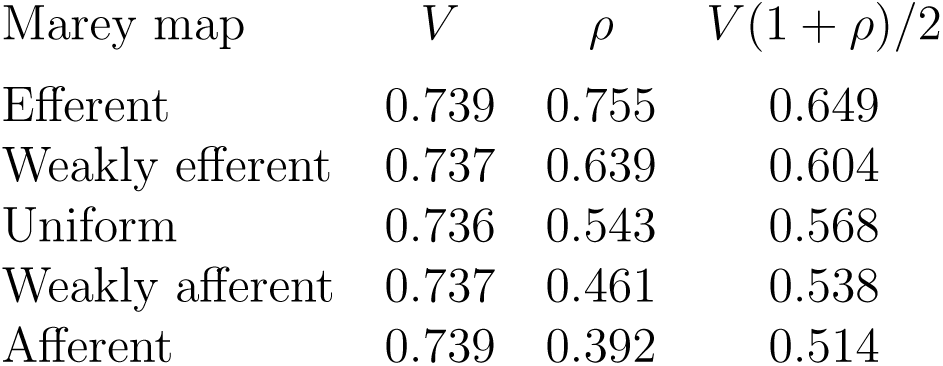
Variance (V) of physical length shared within two half chromosomes, and the correlation (*ρ*) between sharing in one and in the other, for a range of Marey maps (the three examples of Fig. 1 and two intermediate cases). Each segment has map length 0.5M. Variance of sharing for the whole chromosome is V(1 + *ρ*)/2. All values are scaled by the variance of shared map length for a 1M chromosome.

| Marey map | $V$ | $\rho$ | $V(1 + \rho)/2$ |
| --- | --- | --- | --- |
| Efferent | 0.739 | 0.755 | 0.649 |
| Weakly efferent | 0.737 | 0.639 | 0.604 |
| Uniform | 0.736 | 0.543 | 0.568 |
| Weakly afferent | 0.737 | 0.461 | 0.538 |
| Afferent | 0.739 | 0.392 | 0.514 |

### Recombination hotspots

Consider a chromosome with constant recombination rate except for one or more idealised recombination hotspots, each with finite map length but assumed zero physical length. The hotspots divide the chromosome into segments, within each of which the recombination rate is assumed to be constant. The standard broken-stick calculation applies, except that in eqn (7), the exponential terms for intervening segments must also include any intervening hotspots. The effect of a single hotspot placed at different positions on the chromosome is shown in Fig. 2, and the effect of positioning two hotspots is shown in Fig. 3. The variance obtained with a single central hotspot is generally less than if the recombination rate is constant along the chromosome. Thus, taking the case of constant recombination rate as the baseline, a single hotspot reduces variance when it is centrally placed, and increases variance when it is near a chromosome end. The effect (increase or decrease) is generally small, however. The maximum effect of two hotspots occurs when both hotspots are at a chromosome end (either with one at each end or with both at the same end). Similar results to those illustrated in Figs 2 and 3 are seen with a wide range of relationships and map lengths.

**Figure 2:**
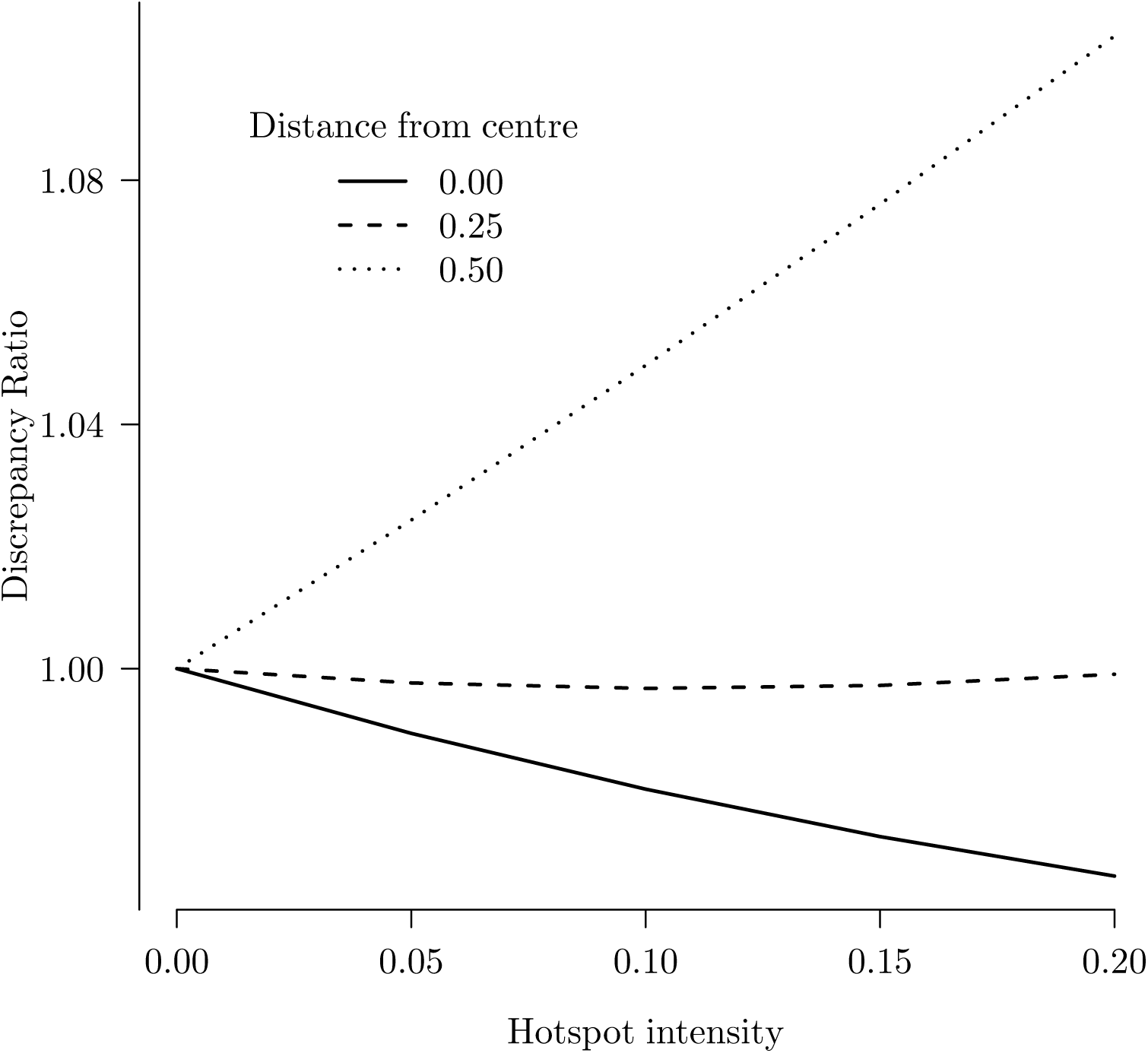
The discrepancy ratio arising from a single hotspot of various intensities placed at different positions on a chromosome of map length 1M. The ‘map length’ of the hotspot is 0.2M and assumed relationship is GPO.

**Figure 3:**
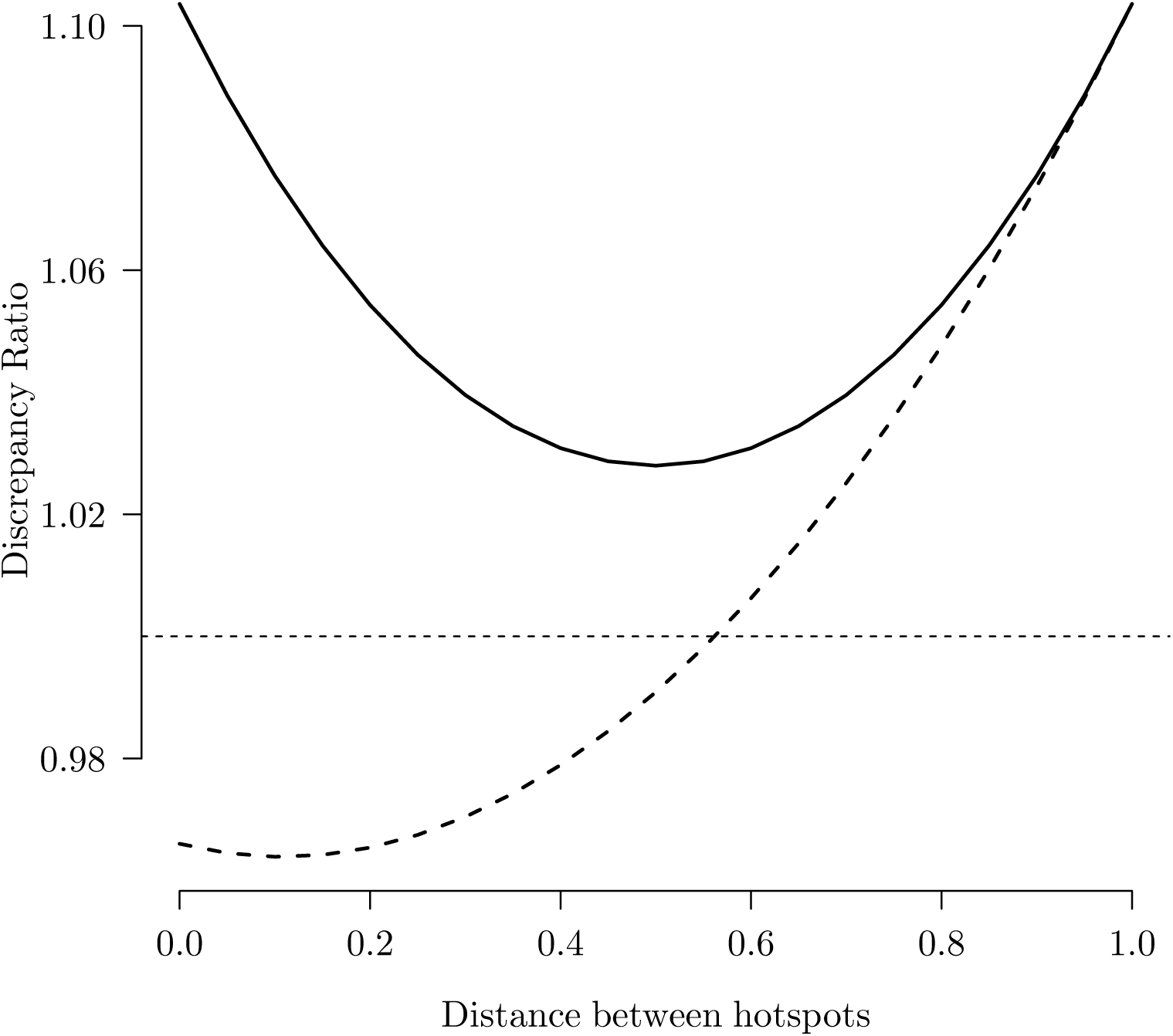
Effect on the discrepancy ratio of different spacings between chromosome ends and two hotspots. A background of constant recombination rate is assumed. The relationship is GPO, the chromosome map length is 1M, and the ‘map length’ of each hotspot is 0.1M. Lower graph: outside spacings are constrained to be equal, and hotspots are symmetrically placed about the chromosome centre. Upper graph: one hotspot is placed at a chromosome end. The area between the two graphs represents varying degrees of asymmetry in hotspot placement.

### Broad-scale variation in recombination rate

We now consider the impact of broad-scale variation in recombination rate. A broken-stick model with three segments of equal physical length but different map lengths was found sufficiently flexible to describe many situations. Let *ℓ*_1_,*ℓ*_2_ and *ℓ*_3_ be the proportional map lengths of the left, middle and right segments, so *ℓ*_1_ + *ℓ*_2_ + *ℓ*_3_ = 1.

For the GPO relationship, and a chromosome with map length 2M, Fig. 4 shows the effect of changing *ℓ*_2_, (a) with *ℓ*_1_ = *ℓ*_3_, and (b) with zero map length for one of the two outside segments. Points between graphs (a) and (b) represent a division of map lengths between outside segments less extreme than case (b). The maximum disparity occurs when all recombination is concentrated in one end segment. This is in agreement with our results for hotspots, but now the magnitude of the effect is much greater.

**Figure 4:**
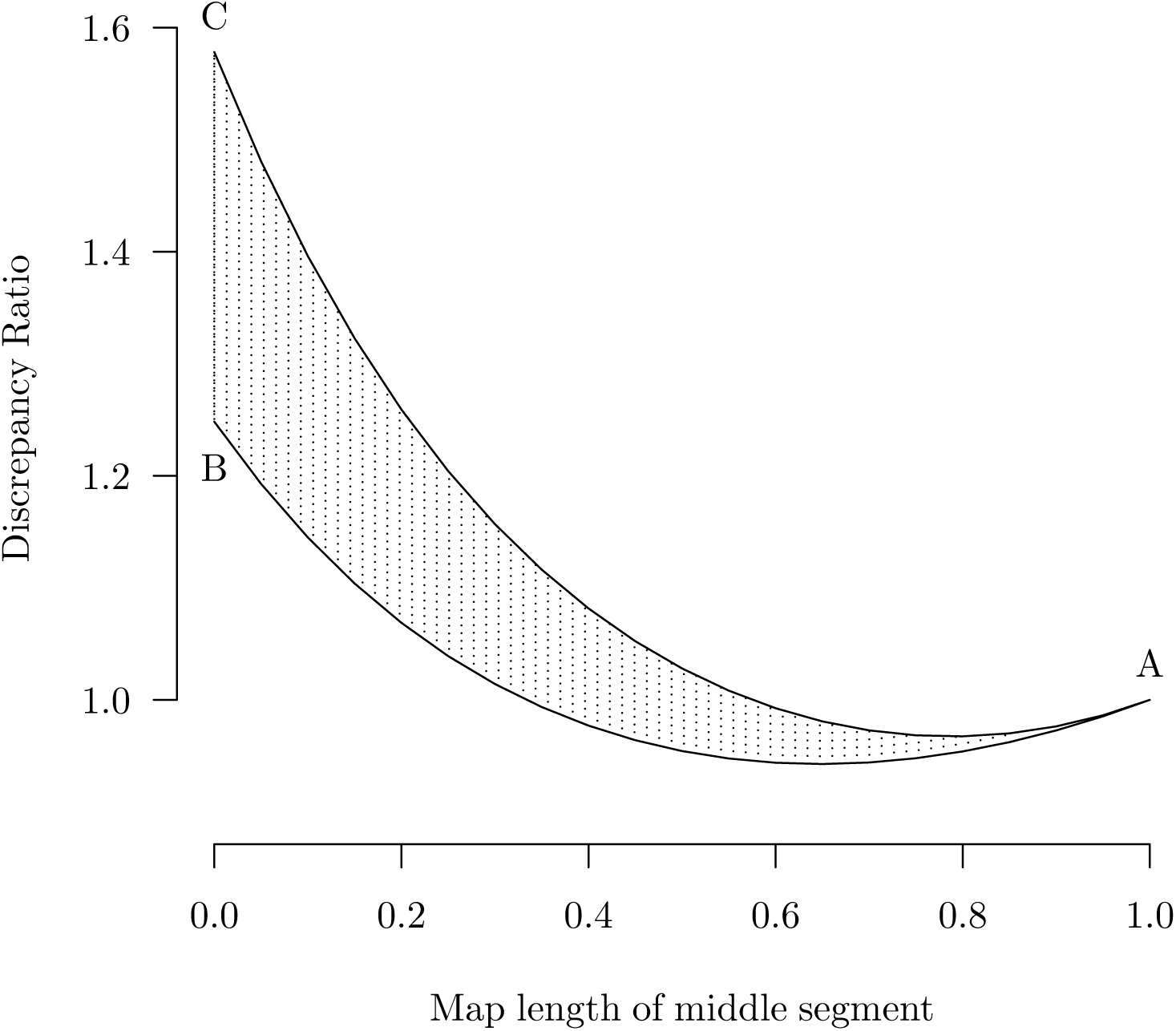
Effect on the discrepancy ratio of the map length (*ℓ*_2_) of the middle segment of a three-segment broken stick, and of different ways of partitioning the remaining map length (*ℓ*_1_ + *ℓ*_3_) between the two ends. GPO relationship, map length 2M. Lower graph: the effect of changing *ℓ*_2_ with the residual map length divided equally between the outside segments (*l*_1_ = *l*_3_); upper graph, the extreme case where all of the residual map length is assigned to one of the outside segments, with no recombination in the other. The area between the two graphs represents the effect of unequal assignment to the outside segments. Three extreme points A, B, and C are discussed in the text.

Results for other relationships and map lengths are expressed in terms of three extreme points, labelled A, B and C in Fig. 4. At point A, all recombination is confined to the middle segment, with end segments either completely sharing or completely non-sharing. At B, there is no recombination in the middle segment, with chromosome map length equally divided between the end segments. At C, all recombination is confined to one of the end segments. For different map lengths and relationships, the area bounded by the three extreme points retains the same basic shape, but A, B and C move up or down in fairly close correlation (Fig. 5). The largest possible variance ratio increases as the relationship becomes more remote. Changes in A, B, and particularly C become more pronounced as map length is increased. Overall, the potential for an increase in variance is greatest with more remote relationships and longer chromosome lengths. Similar trends with map length and bilineal relationship are seen in the variance of double sharing (Fig. 6).

**Figure 5:**
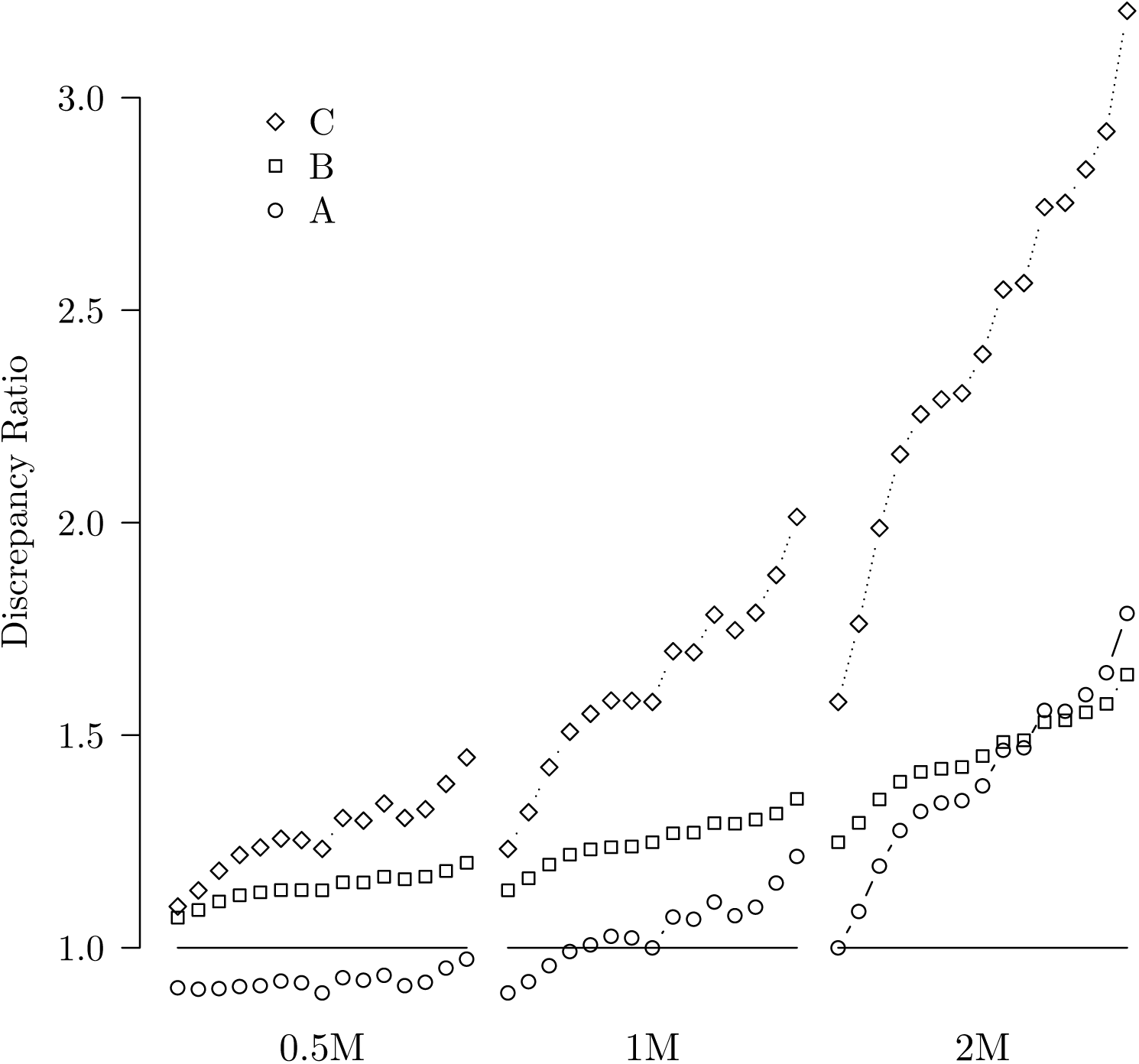
Discrepancy ratios for Marey maps represented by points A, B and C of Fig. 4, for map lengths 0.5M, 1M, and 2M, and a range of relationships: GPO, G2PO, G3PO, HUN, G4PO, GUGN, HC, HS, HC1R, G5PO, C1R, UN, FC, CC, 2C1R, left to right in that order. Values for DHC, DFC, and FS are the same as for HC, FC, and HS, respectively.

**Figure 6:**
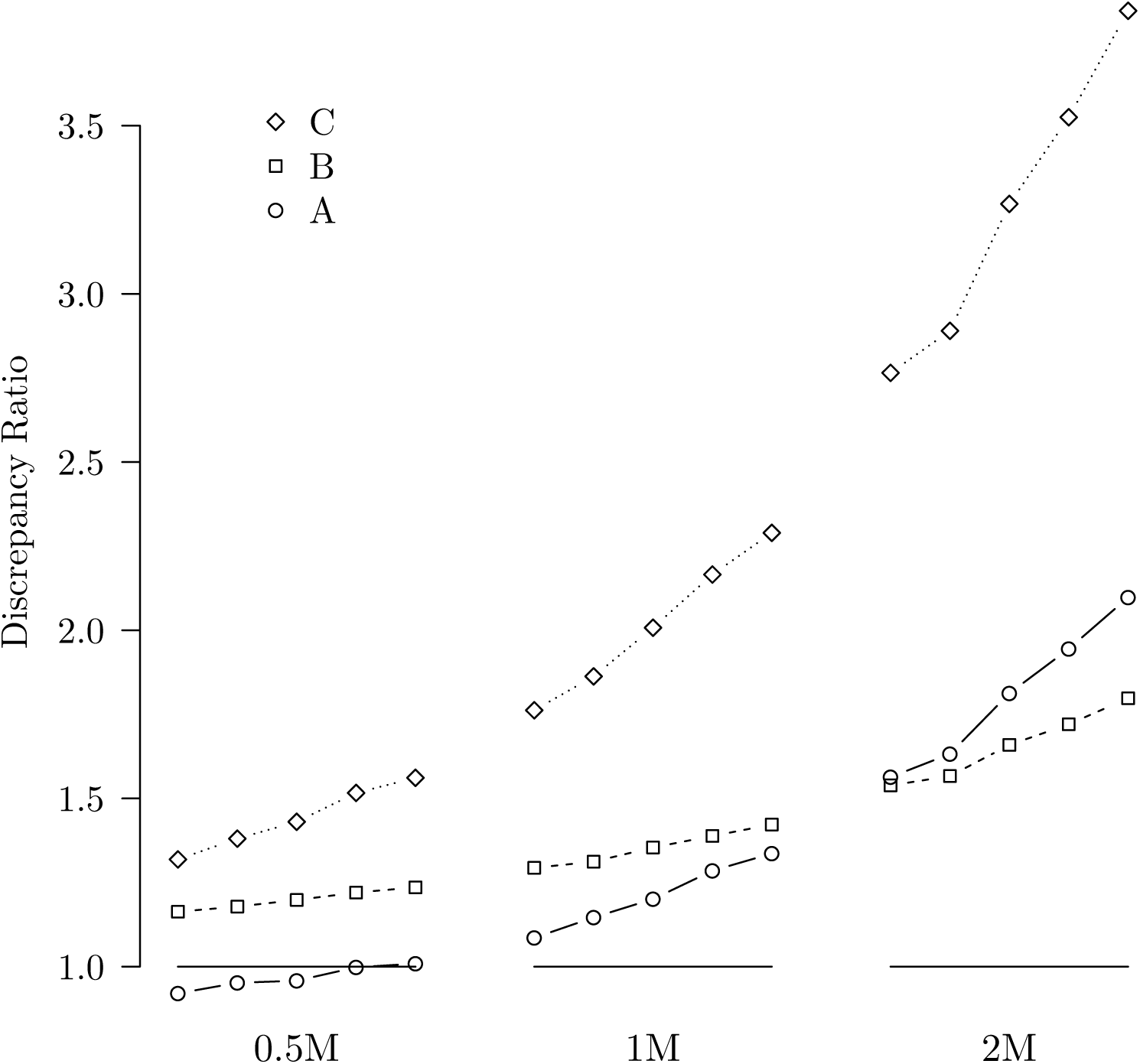
Discrepancy ratios for double sharing for Marey maps represented by points A, B and C of Fig. 4, using map lengths 0.5M, 1M, and 2M, and a range of bilineal relationships: FS, HSHC, FSFC, DHC, DFC, left to right in that order.

### Example: The chicken genome

We investigated variance of identity for the chicken genome. Our data comprised physical and map distances for segments between the 9268 markers in the published linkage map for GGA1-GGA27 (Groenen et al., 2009, Supplemental Table 1). We omitted GGA22 and GGA25, which showed errors in the sequence assembly, as well as GGA16 and the sex chromosomes GGAZ.

Theoretical variances for a GPO relationship were calculated on both map and physical scales, using a broken-stick model for the latter, and the ratio of physical-length variance to map-length variance was plotted separately for each chromosome (Fig. 7). This ratio increases with map length, exceeding 1.0 for the longer chromosomes, less than 1.0 for some of the shorter chromosomes.

**Figure 7:**
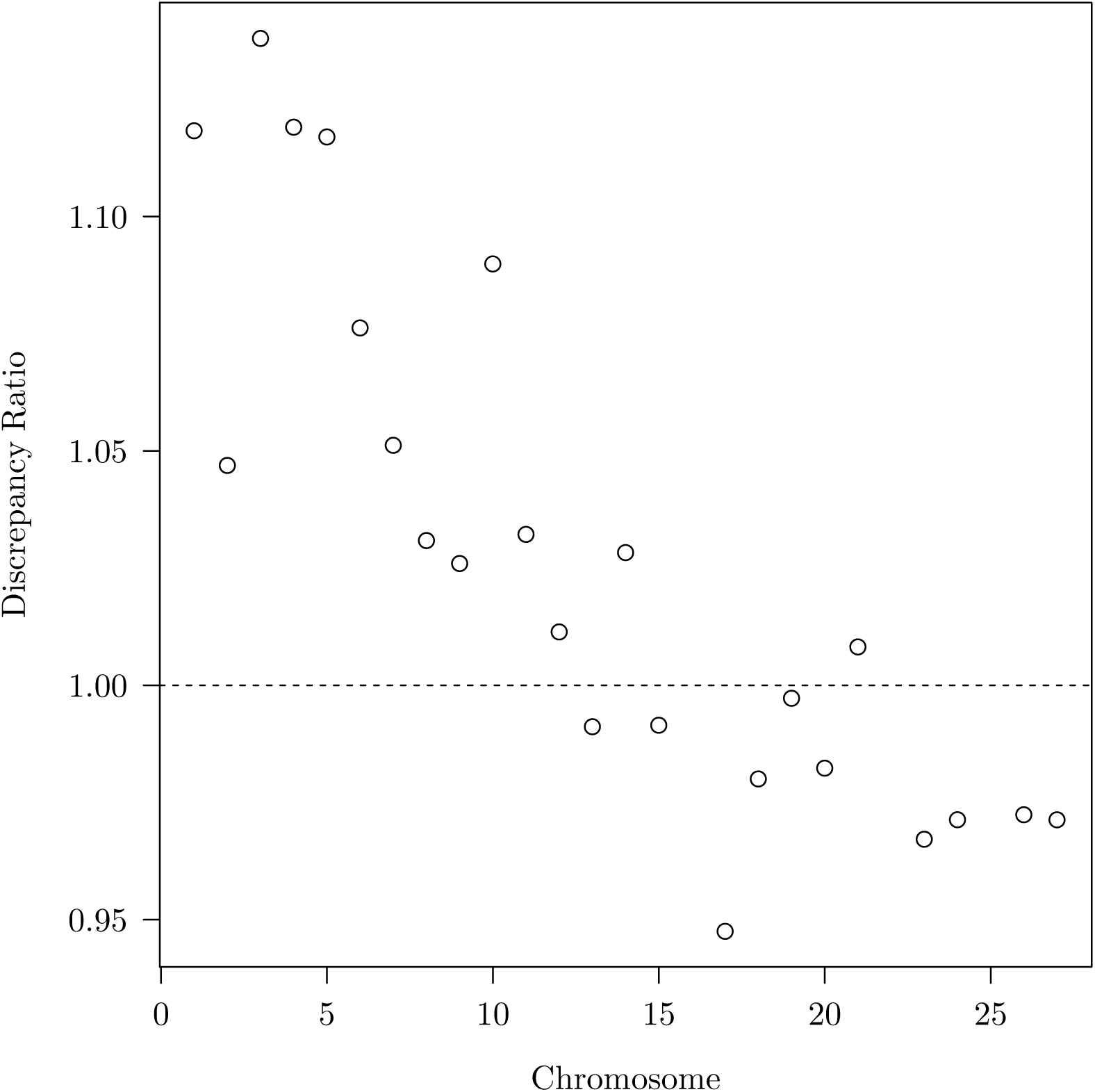
Discrepancy ratios for chicken chromosomes – GGA15, GGA17 – GGA20. GGA16 and chromosomes smaller than GGA20 did not have sufficient data to produce reliable estimates. Assumed relationship GPO.

Similar results are seen with more remote relationships.

### Genome-wide sharing of identity

Our calculations have so far been focussed on a single chromosome. For a genome comprising several independently segregating chromosomes, the overall proportion shared is an average of the values for the individual chromosomes, weighted by their lengths, and the variance of the proportion of genome shared is 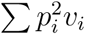, where *p_i_* is the relative length and *v_i_* the variance of sharing of the ith chromosome. The values (*p_i_*, *v_i_*) can be calculated on the scale of map or physical length. For the 24 chicken chromosomes, although the discrepancy ratio for individual chromosomes never exceeded 15% (Fig 7), the variance of the genome-wide proportion of physical length shared was 38% greater than that on the map scale.

## DISCUSSION

We have shown that, when recombination rate varies along the chromosome, variation in relationship as measured in terms of the physical length of the genome can differ quite substantially from that measured in map distance or map units. Among the main influences on the difference are recombination hotspots and cases where much of the recombination generated by meiosis is found towards one chromosome end. Nevertheless, the effect is likely to be less than that of the total map length of the genome, the number of chromosomes, or their map lengths. For example, the chicken genome shows substantial heterogeneity in recombination rate both between and within chromosomes (Groenen et al., 2009), but heterogeneity in rate between chromosomes does not affect variance of sharing. The longest chromosomes have a nearly constant rate except at the chromosome ends where the influence of the higher recombination rate is likely to be small because a small amount of genome is involved. The shorter chromosomes tend to have higher mean recombination values than do the long chromosomes and so contribute rather little to the overall variance in relationship. Some however, show strong differences in recombination rate between the ends.

In general, proportional changes in variance (compared to the reference case) are similar for a wide range of relationships, tending to increase as relationships become more distant (and absolute values decrease). They are greatest with high recombination rates near chromosome ends, and increase with map length. In view of the wide range investigated, degree of relationship has a rather small impact. Clearly the contribution of the higher decay functions which affect only the more distant relationships (Table 2) are not making a major contribution. Effects of map length and non-linearity in the Marey map are much greater.

If variance in relationship can differ depending on whether it is based on physical or map length, this raises the question of which is the more appropriate quantity to analyse. Map length does not necessarily reflect parts of the genome with high gene density but low recombination rate, for example. In some applications, map length may be the appropriate quantity because it is potentially the best indicator of local linkage disequilibrium. This is relevant when using genomic data to predict genetic merit in livestock and crops (Meuwissen et al., 2001) or complex human diseases (Purcell et al., 2009). Here the issue is how much information about the genotype at trait loci comes from neighboring markers, and marker linkage is likely to be the most suitable quantity with which to weight observations.

Although we have shown that it is theoretically possible for the impact of differences between physical and map distances (apart from scale) to be substantial, in practice the impact seems to be limited. Compared to the vast range of recombination rates among different orders of animals and even species, the effects on variation in relationship are relatively small. In *Drosophila melanogaster,* for example, there is no recombination in males, and most of the genome is on three chromosomes. In birds, physical and map lengths are highly heterogeneous, and within chromosomes there can be areas of extremely high recombination rate. Therefore it seems reasonable to assume that the chromosome architecture is not subject to very strong evolutionary pressure beyond quite crude limits, for example a basic need for telomeres and perhaps a centromere. There are selective forces that are likely to increase recombination rates, for example by increasing the probability of survival of favorable mutants tightly linked to regions under selection, or by increasing retention of epistatic complexes in the face of recombination. However, it is difficult to imagine selective forces sufficiently divergent to explain the wide variations observed in genomic structure.

## APPENDIX 1

### The ibd process

Let *X* (*t*) be a random variable which takes the value 1 when there is sharing at locus *t* and is otherwise zero. Equations (1) and (2) are respectively *E*{*X*(*s*)*X*(*t*)} and cov{*X*(*s*), *X*(*t*)} for loci at *s* and *t* separated by map distance *d*. For all *t*, *E*(*X* (*t*)) = *k* is the probability of sharing at locus *t*. If, instead of regarding *s* and *t* as indexing discrete chromosomal segments, we assume that the number of loci are effectively infinite, *s* and *t* become continously varying parameters identifying point locations on the chromosome, and the proportion of chromosome shared is the ‘time’ average

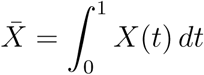

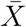 is a random variable (different realisations of the process *X* (*t*), 0 < *t* < 1 give rise to different values of 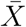), and 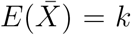. The variance of sharing, or identity, is the average covariance

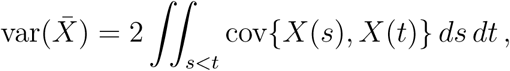

which is eqn 3. The positions *s* and *t* are relative to chromosome ends, and can represent either physical or map distances.

### The Markov chain representation

An alternative derivation of equation (1) and the coefficients in Table 2 is obtained by regarding the ibd process as a continuous-time Markov chain with the ‘time’ axis representing position on the chromosome (Donnelly, 1983; Guo, 1995). We give two examples.

The GPO relationship is represented by a chain with two states, one of which represents sharing. The probability of being in the sharing state at time *t*, given that the chain starts in the sharing state, is (1/2) [1+exp(−2*t*)], showing how the chain reverts to equilibrium after starting in the ‘sharing’ state. The probability of sharing at *t* = 0 and again at time *t* is (1/2)^2^[1 + exp(−2*t*)] which is the probability of simultaneous sharing at two linked loci given by eqn (1) for the GPO relationship.

For the G2PO relationship, the chain has four states, one of which represents sharing, and the conditional probability of sharing at time *t*, given sharing at *t* = 0, is

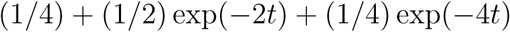

Here, the return to equilibrium is more complex, with *p*_2_(*t*) = *p*_3_(*t*) for all *t, p*_1_(*t*) −*p*_4_(*t*) = exp(−2_t_), and *p*_1_(*t*) −*p*_2_(*t*) −*p*_3_(*t*) +*p*_4_(*t*) = exp(−4*t*), where *p_i_*(*t*) is the probabilility of being in state *i* at time *t,* given that the chain starts in the sharing state (state 1). The probability of sharing at *t* = 0 and again at time *t* is

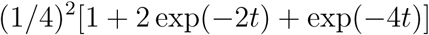

which can be compared with eqn (1) and the entry for G2PO in Table 2.

## LITERATURE CITED

Backström, N., Forstmeier, W., Schielzeth, H., Mellenius, H., Kiwoong, N., et al. (2010). The recombination landscape of the zebra finch *Taeniopygia guttata* genome. Genome Res., 20:485–495.

Chakravarti, A. (1991). A graphical representation of genetic and physical maps: the Marey map. Genomics, 11:219–222.

Charlesworth, B. and Charlesworth, D. (2010). Elements of Evolutionary Genetics. Robertson & Co., Colorado.

Donnelly, K. P. (1983). The probability that related individuals share some section of the genome identical by descent. Theor. Pop. Biol., 23:34–63.

Franklin, I. R. (1977). The distribution of the proportion of the genome which is homozygous by descent in inbred individuals. Theor. Pop. Biol., 11:60–80.

Groenen, M. A. M., Wahlberg, P., Foglio, M., Cheng, H. H., Megens, H.-J., et al. (2009). A high-density SNP-based linkage map of the chicken genome reveals sequence features correlated with recombination rate. Genome Res., 19:510–519.

Guo, S.-W. (1995). Proportion of genome shared identical by descent by relatives: concept, computation, and applications. Am. J. Hum. Genet., 56:1468–1476.

Haldane, J. B. S. (1919). The combination of linkage values, and the calculation of distance between linked factors. J. Genet., 8:299–309.

Hill, W. G. (1993). Variation in genetic composition in backcrossing programs. J. Hered., 84(3):212–223.

Hill, W. G. and Weir, B. S. (2011). Variation in actual relationship as a consequence of Mendelian sampling and linkage. Genet. Res., 93:47–74.

Hill, W. G. and Weir, B. S. (2012). Variation in actual relationship among descendants of inbred individuals. Genet. Res., 94:267–274.

Kong, X., Murphy, K., Raj, T., He, C., White, P. S., and Matise, T. C. (2004). A combined physical-linkage map of the human genome. Am. J. Hum. Genet., 75:1143–1148.

Matise, T. C., Chen, F., Chen, W., De La Vega, F. M., Hansen, M., et al. (2007). A second-generation combined linkage-physical map of the human genome. Genome Res., 17(12):1783–1786.

McVean, G. A. T., Myers, S. R., Hunt, S., Deloukas, P., Bentley, D. R., et al. (2004). The fine-scale structure of recombination rate variation in the human genome. Science, 304:581–4.

Meuwissen, T. H. E., Hayes, B. J., and Goddard, M. E. (2001). Prediction of total genetic value using genome-wide dense marker maps. Genetics, 157:1819–1829.

Purcell, S. M. et al. (2009). Common polygenic variation contributes to risk of schizophrenia and bipolar disorder. Nature, 460:748–752.

Ritz, K. R., Noor, M. A. F., and Singh, N. D. (2017). Variation in recombination rate: Adaptive or not? Trends in Genetics, 33:364–374.

Schwartz, J. J., Roach, D. J., Thomas, J. H., and Shendure, J. (2014). Primate evolution of the recombination regulator PRDM9. Nat. Comm., 5:4370.

Stam, P. and Zeven, A. C. (1981). The theoretical proportion of the genome identical by descent in finite populations. Genet. Res., 35:131–155.

Stapley, J., Feulner, P. G. D., Johnston, S. E., Santure, A. W., and Smadja, C. M. (2017). Variation in recombination frequency and distribution across eukaryotes: patterns and processes. Phil. Trans. Roy. Soc. B, 372:20160455.

Visscher, P. M. (2009). Whole genome approaches to quantitative genetics. Genetica, 136:351–358.

Wright, S. (1922). Coefficients of inbreeding and relationship. Am. Nat., 56:330–338.

